# Tracing the History of LINE and SINE Extinction in Sigmodontine Rodents

**DOI:** 10.1101/242636

**Authors:** Lei Yang, Holly A Wichman

**Affiliations:** Department of Biological Sciences & Institute for Bioinformatics and Evolutionary Studies, University of Idaho, Moscow, Idaho, United States of America; Present address: Department of Biology, Pennsylvania State University, University Park, Pennsylvania, United States of America

## Abstract

**Background:** L1 retrotransposons have co-evolved with their mammalian hosts for the entire history of mammals and currently make up to 20% of a typical mammalian genome. B1 retrotransposons are dependent on L1 for retrotransposition and span the evolutionary history of rodents since their radiation. L1s were found to have lost their activity in a group of South American rodents, the Sigmodontinae, and B1 inactivation preceded the extinction of L1 in the same group. Consequently, a basal group of sigmodontines have active L1s but inactive B1s and a derived clade have both inactive L1s and B1s. It has been suggested that B1s became extinct during a long period of L1 quiescence and that L1s subsequently reemerged in the basal group.

**Results:** Here we investigate the evolutionary histories of L1 and B1 in the sigmodontine rodents and show that L1 activity continued until after the split of the L1-extinct clade and the basal group. After the split, L1s had a small burst of activity in the former group, followed by extinction. In the basal group, activity was initially low but was followed by a dramatic increase in L1 activity. We found the last wave of B1s retrotransposition was large and probably preceded the split between the two rodent clades.

**Conclusions:** Given that L1s had been steadily retrotransposing during the time corresponding to B1 extinction and that the burst of B1 activity preceding B1 extinction was large, we conclude that B1 extinction was not a result of L1 quiescence. Rather, the burst of B1 activity may have contributed to L1 extinction both by competition with L1 and by putting strong selective pressure on the host to control retrotransposition.

## Background

LINEs (Long INterspersed Elements) are autonomous non-LTR (non-long terminal repeat) retrotransposons that move through an RNA intermediate. L1 (LINE-1) is the most successful family of LINEs in eutherian mammals [1] and make up ~20% of a typical mammalian genome [2, 3]. A functional full-length L1 is typically 6,000-7,000 bp long and composed of a 5’ untranslated region (5’UTR) harboring an RNA polymerase II promoter, two non-overlapping open reading frames (ORFs) known as ORF1 and ORF2 and a 3’UTR followed by a poly-adenosine sequence [4]. The structure of L1 can be diverse among different mammals, particularly in the 5′ UTR and ORF1 [5]. The ORF-encoded proteins are strictly required for L1 retrotransposition and are highly *cis*-preferential [6, 7]. L1s are adenosine rich (~40%) on their coding strand, which results in biased codon usage compared to host genes [8, 9], elongation defects [10], and premature RNA splicing [11]. This A-richness contributes to the inefficiency of L1 retrotransposition and is proposed to regulate the genes in their vicinity [10].

SINEs (Short INterspersed Elements) are relatively short non-autonomous, non-LTR transposable elements. SINEs do not encode proteins for their own retrotransposition and depend on the reverse transcriptase encoded by other transposable elements such as LINEs [12, 13]. Although L1s are highly *cis*-preferential [6, 7], SINEs can take advantage of L1-encoded proteins for their own retrotransposition [12-14]. Despite their short length, SINEs account for ~10% of a typical mammalian genome due to their high copy numbers [2, 3]. Among the ~70 SINE families found in mammals [15], B1 is the most abundant in mouse [3] and possibly most rodent species [16], occupying ~3% of the mouse genome [3]. B1s derived from the RNA component of signal recognition particle 7SL RNA [17, 18] and share features with its ancestors – a functional B1 is ~150 bp long and transcribed by RNA polymerase III with the aid of its two transcription factor binding boxes [19, 20]. B1 sequences are rich in CpG sites, which are methylated and thus prone to mutation in mammalian genomes [21], and the elevated mutation rate is pronounced compared to the A-rich L1s. Because the majority of new L1 and B1 inserts are neutrally-evolving pseudogenes, the CpG-rich B1 sequences decay faster than the A-rich L1 sequences.

Both L1 and B1 have long histories of co-evolution with their host genomes. Unlike some transposable elements, there is no known targeted mechanism for L1s excision and thus L1s persist in the genome unless they are removed by non-specific mechanisms. The oldest L1s trace back to the common ancestor of placental mammals and marsupials, ~160 MYA [1, 22]. L1s evolve as master lineages so that a single or a few lineages are responsible for the total retrotransposition in a short time window [23-26]. New master elements replace the old ones, eventually dominating retrotransposition, and this replacement process happens recurrently. B1s are younger than L1s, having arisen just before the divergence of the common ancestor of rodents, ~65 MYA [27], and they are specific to rodents. Other SINEs, including B2, B4 and ID elements, are also present in rodent genomes [16]. SINE families have been interacting with L1s for more than 100 MYA, and fossil remnants of extinct SINE families are detectable in well-characterized mammalian genomes [15, 28]. Despite being under strict regulation, L1 and B1 make up approximately a quarter of a typical rodent genome [3]. For example, in the mouse genome, there are ~599,000 total copies of L1, responsible for ~19% of the genome [3], of which ~3,000 copies are potentially functional [29], and ~564,000 copies of B1s, responsible for ~3% of the genome [3].

LINEs and SINEs have considerable impact on the mammalian genome, although they were traditionally viewed as “junk DNA”. As LINEs and SINEs, including L1s and B1s, retrotranspose and recombine, they introduce genome instability [30], cause disease [31] and may occasionally be co-opted by the host to serve certain functions, such as their proposed roles in neuro-plasticity [32, 33], X chromosome inactivation [34, 35], regulatory functions [36, 37], DNA break-repair [38] and genome organization [39, 40]. Due to the deleterious effects of LINEs and SINEs on the genome, the hosts have evolved many mechanisms to defend against them [41-45]. In addition, the fact that L1 doesn’t encode all the enzymatic components required for retrotransposition could result in ongoing competition between L1s and the host for these required host factors [46, 47]. Host defense against L1s and B1s are especially strong in germline cells due to germline-specific host defense mechanisms, so that only a limited number of new copies are inserted in each generation [48, 49]. L1s and B1s are both epigenetically silenced [50, 51] and under the control of small RNAs [52], which are specifically expressed in germline cells.

Since L1 retrotransposition is under strict control by multiple host defenses, it might seem reasonable for the host to occasionally win the evolutionary arms race with L1s, resulting in loss of L1 activity (L1 extinction). L1s are not known to move horizontally, so such extinctions would affect all derived host species. Two factors are of note here. First, clades with early L1 extinctions could have given rise to large mammalian lineages without L1 activity and be easily detected because of both the number of species affected and the deterioration of the remnant sequences in the genome. Secondly, recent extinctions will be difficult to differentiate from periods of L1 quiescence. To clarify the terms related to loss of L1 activity in this work, we refer to a period of low L1 activity as “quiescence” and complete loss of L1 activity as “extinction”. Given the large phylogenetic impact of early extinctions, one might expect L1s to eventually become extinct in most mammalian genomes, and yet L1s have persisted throughout the entire evolutionary history of their placental mammal and marsupial hosts. Thus, either most L1 extinctions are either recent or rare, or mammalian lineages subject to ancient L1 extinctions do not persist or they give rise to few new species. Understanding the dynamics of L1 extinction will be as important as understanding the dynamics of L1 activity in sorting out the impact of L1s on mammalian genome evolution.

Several cases of L1 extinction have been proposed in the literature [53-61] and two of these are deep extinction events that cover major groups of mammals [53-57]. One of the major L1 extinctions [55-57] occurred in a large group of South American rodents and includes most species in Sigmodontinae. Sigmodontinae is a subfamily of the Cricetidae family, including approximately 377 species classified into 74 genera in nine tribes (Figure 1) [62] and thus contains to 7-8% of the estimated 5,000 mammalian species [63]. Given that B1 retrotransposition is dependent on that of L1, it is expected that B1s should lose their activity simultaneously with L1s. However, the B1 extinction in Sigmodontinae appears to have preceded that of L1s based on samples from 14 genera in five tribes [55-57], where the basal genus *Sigmodon* carries inactive B1 and active L1, and the descendant genera carry both inactive L1 and B1 (Figure 1). It has also been shown that loss of L1 and B1 activity follows the expansion of a group of endogenous retrovirus [64, 65].

**Figure 1.**
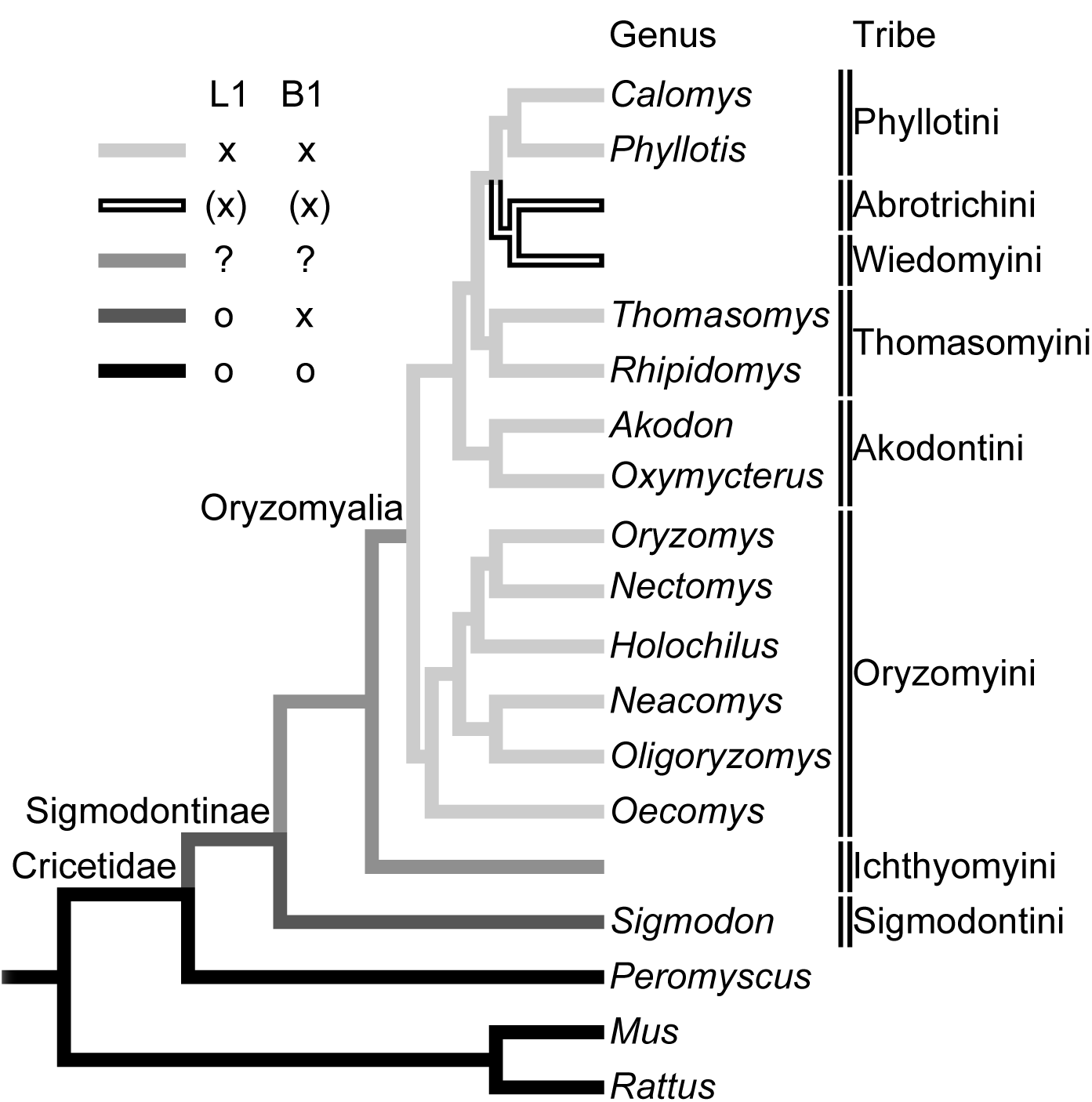
The phylogeny of the sigmodontine rodents. The tree is based on Schenck *et al.* [68]. Taxa are the sampled genera in the group; tribes are indicated on the right side of the taxa. Eight of the nine tribes and 12 of the 14 sampled genera by Rinehart *et al.* [57] are shown. L1 and B1 activity of each taxon is demonstrated by gray scale and: black indicates active L1 and B1, dark gray indicates active L1 and inactive B1 and medium gray indicates the taxa where L1 activity cannot be inferred and light gray indicates the taxa where L1 can be inferred to be active. “o” corresponds to active L1 and B1 and “x” corresponds to inactive L1 and B1.

It was previously hypothesized by Cordaux and Batzer that the L1 can experience long-term quiescence as a “stealth driver” [66], and B1 extinction could have happened during this period of L1 quiescence [57]. Since B1s are more prone to mutations than the average sequence due to enriched CpG content, Rinehart *et al*. [57] hypothesized that B1 was unable to retrotranspose at a high enough rate during L1 quiescence to replace their active copies, accumulating debilitating mutations more rapidly [21] than L1s. When a more active family of L1 emerged in the Sigmodontini, B1 was too degenerated to retrotranspose, resulting in B1 extinction even in the presence of high L1 activity.

In this study, we investigate the evolution histories of L1 and B1 spanning the time of their extinctions and the radiation of the extant species in Sigmodontinae (Figure 1). Since the group carrying extinct L1s and B1s (Oryzomyalia, Figure 1) shares a common ancestor, we used the marsh rice rat *Oryzomys palustris* to represent this group, hereafter referred to as the “L1-extinct clade”. We used the hispid cotton rat *Sigmodon hispidus* to represent the clade carrying active L1 but inactive B1, hereafter referred to as the “basal group”. We used the deer mouse *Peromyscus maniculatus* to represent a closely related clade carrying both active L1 and B1, hereafter referred to as the “outgroup”.

Using genome trace files from the species representing the L1-extinct clade and the basal group, we show that the activity of L1 and B1 families that precede the divergence of the clades is comparable in the current genomes of the two groups. L1 families had been steadily replaced before the split of the two groups and maintained activity after the split of the basal group and the L1-extinct clade. Shortly after this split L1 activity ceased in the L1-extinct clade but became highly active in the basal group. B1s, on the other hand, had a very large increase in activity prior to the split between the L1-extinct clade and the basal group, and there is no strong evidence of activity in the two groups following their divergence. The large burst of B1 activity just prior to extinction suggests that L1 quiescence is unlikely responsible for B1 extinction. The last wave of B1 retrotransposition is the largest detectable in the B1 evolutionary history of the group, suggesting B1s’ strong competition with L1s or enhanced host defense triggered by radical B1 expansion might have contributed to the extinction of L1.

## Results

To investigate the history of L1 retrotransposition in *O. palustris* and *S. hispidus*, we used COSEG [67] to identify closely related L1 groups based on shared, co-segregating sites as described in Methods. We follow the convention of COSEG to designate these groups as *subfamilies*. RepeatMasker [67] was used to initially assign genomic L1 copies to subfamilies, and seven subfamilies with no assigned sequences were removed from further consideration, leaving 47 subfamilies for further analysis.

To examine the activity of L1s in *O. palustris* and *S. hispidus*, we searched the trace files of both genomes separately with the consensus sequences of the abovementioned 47 subfamilies and identified 19,254 sequences in *O. palustris* and 90,526 in *S. hispidus*. The age of each sequence was approximated by its percent divergence from the corresponding subfamily consensus – the higher the percent divergence, the older the sequence. The peak of the distribution was used as an approximation of the age of the subfamily (Table S1). Given the possible changes of evolution rate in the detectable range of L1 evolutionary, a global conversion from percent divergence to time is challenging. However, because of the shared evolutionary history of *O. palustris* and *S. hispidus*, percent divergence is a reasonably good marker to compare the age of L1 subfamilies of the two species.

Subfamily consensus sequences were also subjected to phylogenetic analysis (Figure S1). Subsequently, phylogenetic relationships and sequence similarities between subfamilies were used to assign subfamilies to families with the stipulation that the pairwise distance between subfamilies within a family be no greater than 3.5%. This distance was determined operationally based on the divergences among phylogenetically clustered subfamilies. Clusters of subfamilies that were similar at the sequence level but differed in age were assigned to different families. This process identified five families specific to *S. hispidus* (S1 to S5), four families shared by *O. palustris* and *S. hispidus* (OS1 to OS4) and two shared by *P. maniculatus*, *O. palustris* and *S. hispidus* (OSP1 and OSP2, Table S1). A distance-based phylogeny reflecting the relationship between L1 families is presented in Figure 2A. Individual sequences were assigned to the families to which their subfamilies belong; the age distribution within a family is based on the distance of each sequence from its subfamily consensus (Figure 3).

**Figure 2.**
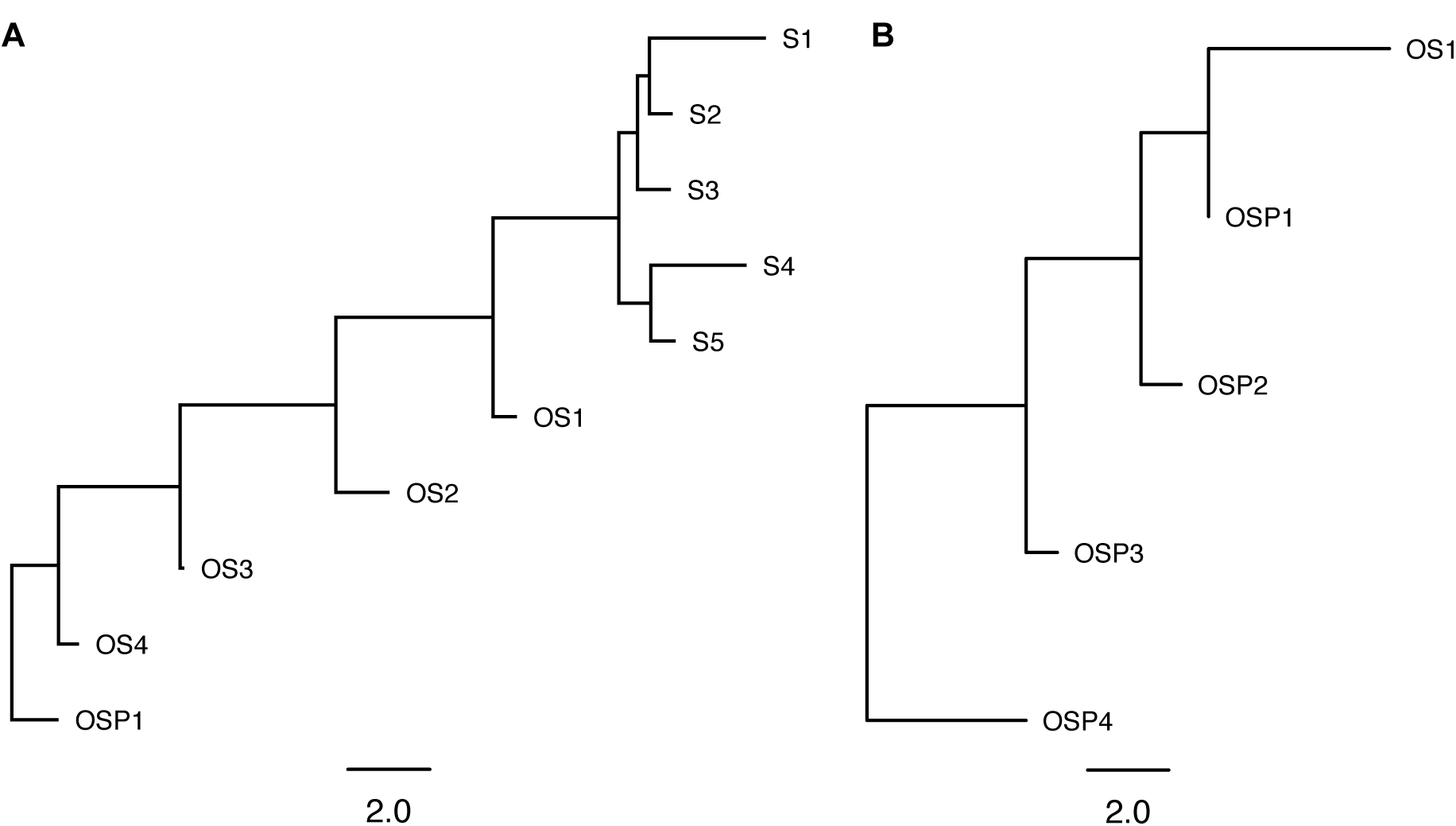
The phylogenies of L1 and B1 families. Panel A shows the L1 tree and B shows the B1 tree. To reflect ages of the families, the trees were based on the distance between families. The distance between any two families was calculated by taking the average pairwise distance of the consensus sequences of subfamilies that belong to each family.

**Figure 3.**
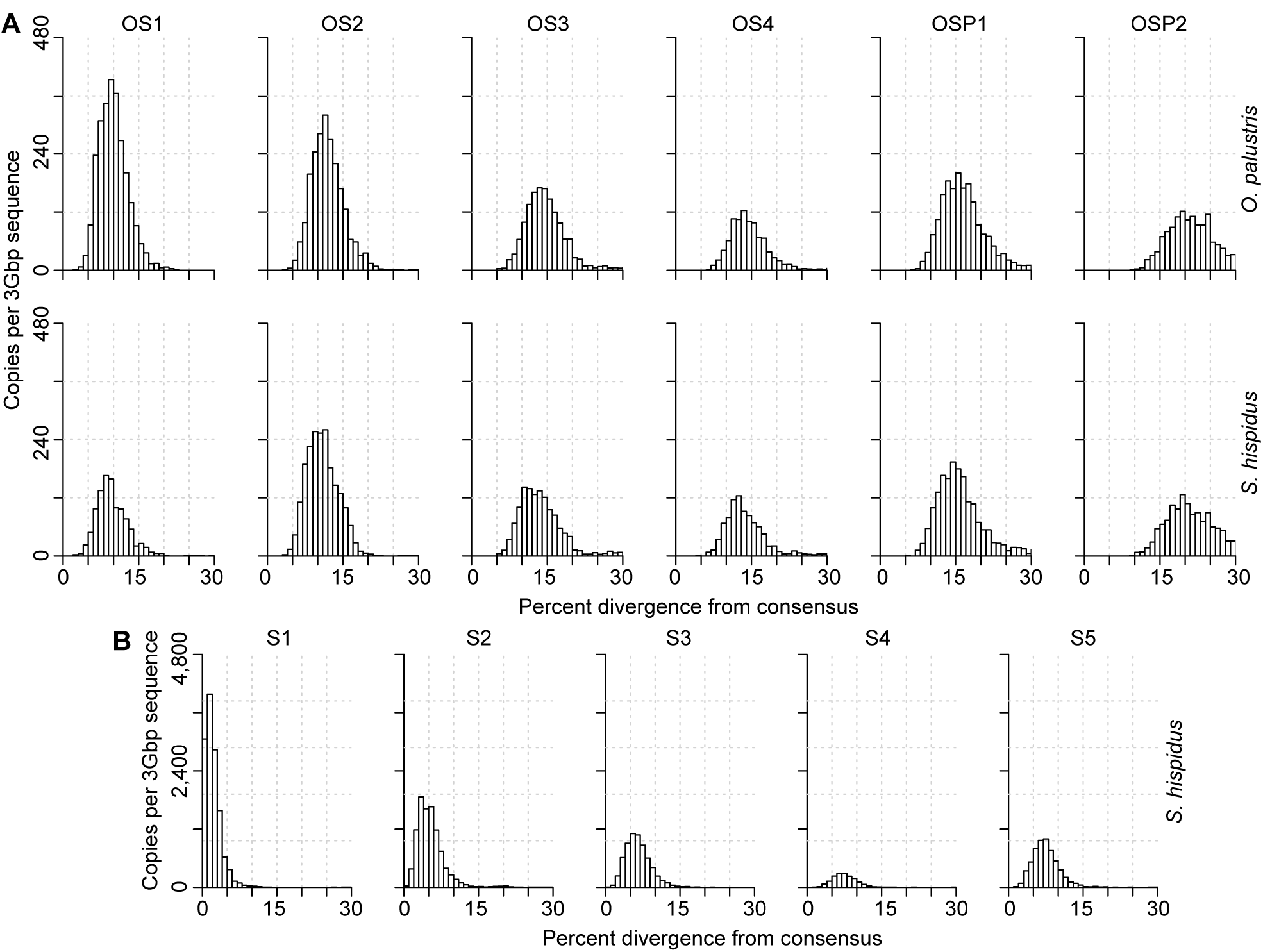
The age distribution of L1 families. L1 families in each row are arranged in chronological order with the youngest families on the left. The species analyzed in each row is indicated at the right. Names of families are noted on the top of each panel. L1 copy number is plotted by percent divergence from the corresponding subfamily consensus in 1% windows. The age of each family is approximated by the peak of the distribution. L1 copy numbers are normalized as copies per three Gbp of MiSeq sequence which approximates the copy number per haploid genome. Panel A shows the shared families and panel B shows the *Sigmodon*-specific families.

As expected, sequences from L1 families shared by *O. palustris* and *S. hispidus* are present in both genomes, and these shared families are fairly synchronized in time and comparable in copy number (Figure 3A). The *Sigmodon*-specific L1 families (Figure 3B, families S1-5) experienced substantial amplification after divergence from the L1-extinct clade, whereas no *Oryzomys*-specific subfamilies were identified by COSEG. The *Sigmodon*-specific subfamilies had a few sequences from the *O. palustris* genome assigned to them, but these assignments appear to be anomalous since the sequences are highly divergent from the subfamily consensus sequences (Table S1). Family OS1, the youngest shared family is of special interest. Family OS1 corresponds to a single L1 subfamily, suggesting that there was little divergence of L1s within the family. It is the last active family prior to the L1 extinction and has ~1.5-fold higher copy numbers per Gbp of sequence in *O. palustris* than in *S. hispidus*. This difference in L1 deposition between *O. palustris* and *S. hispidus* suggests that L1s remained active in the L1-extinct clade after the separation of that group from the basal group. Furthermore, L1s were more active in the lineage leading to Oryzomyalia, in which L1s eventually became extinct, than in the lineage leading to Sigmodontini. A direct comparison of the activity of the L1 families directly preceding this split (OS2), directly following the split (OS1) and at the base of the Sigmodontini (S5) is presented in Figure 4A. Thus, L1 experienced an expansion (family OS1) in the lineage leading to Oryzomyalia immediately before L1 extinction, while the lineage leading to Sigmodontini experienced a delayed but much larger L1 expansion.

**Figure 4.**
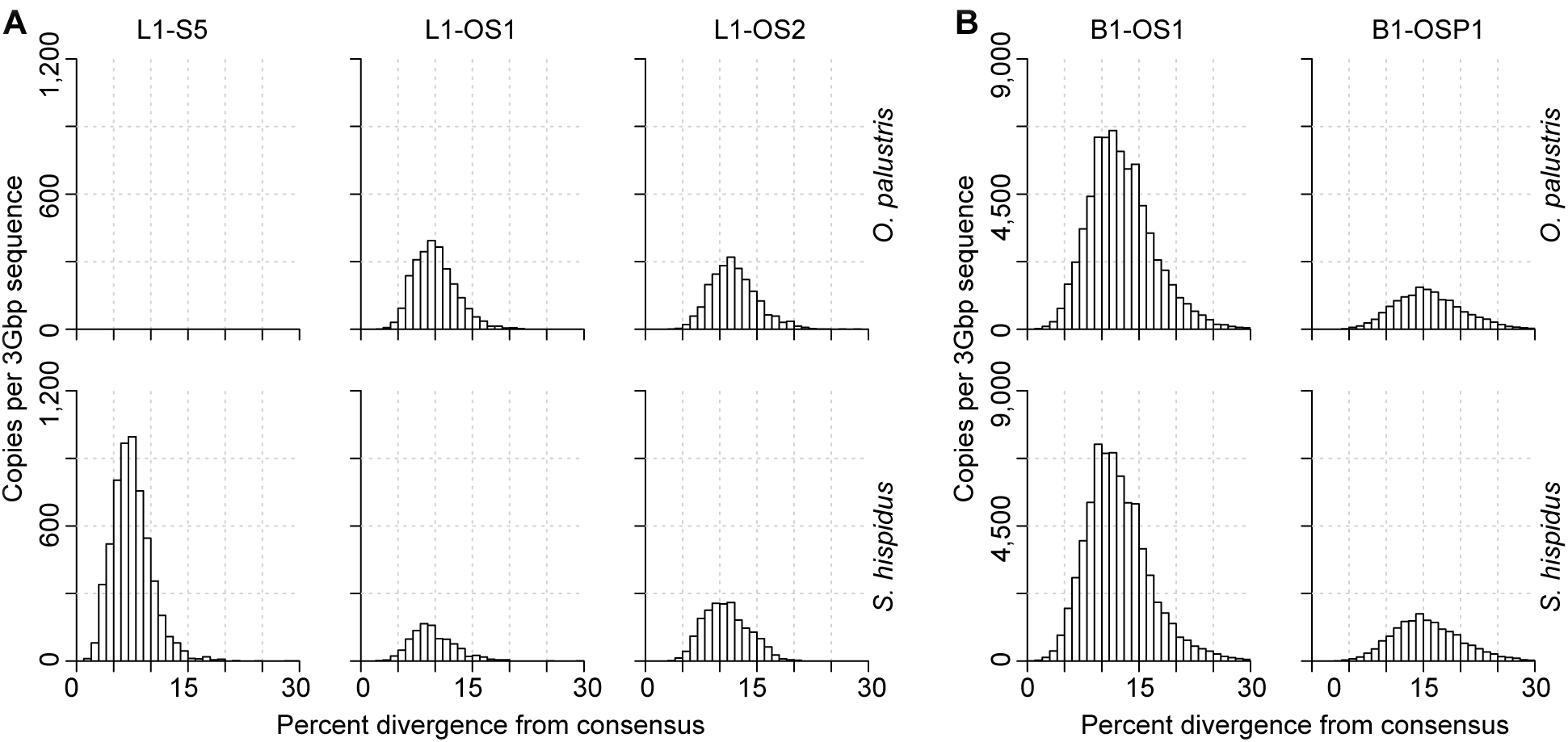
Comparison of L1 and B1 families spanning their extinction. Panel A presents L1 families S5, OS1 and OS2 arranged in a chronological order with the youngest families on the left, and panel B presents B1 families OS1 and OSP1. The species analyzed in each row is indicated at the right. Names of families are noted at the top. Copy number of L1 OS2 is comparable in *O. palustris* and *S. hispidus*, but more OS1 copies were detected in *O. palustris*. Subsequently, there was a new wave of L1 retrotransposition in *S. hispidus* (family S5), but no younger waves of L1 retrotransposition events were identified in *O. palustris*. B1 OS1 corresponds to L1 OS2 in terms of age.

In order to study the B1 dynamics in sigmodontine rodents, we performed the analysis on B1 similar to that done on L1. Because of the short length and CpG-rich nature of B1, we required twice as many sequences to form a subfamily in the second round COSEG as described in Methods. The analysis revealed 30 subfamilies and five families of B1 in both species (Table S2). A distance-based phylogeny reflecting the relationships between B1 families is presented in Figure 2B. One of the families (OS1) is shared by *O. palustris* and *S. hispidus* and the other four (families OSP1-5) are shared by *O. palustris*, *S. hispidus* and *P. maniculatus*. All of the B1 families are shared by *O. palustris* and *S. hispidus* and the representation of these families in both genomes is fairly synchronized in time and comparable in copy number (Figure 5). Since the outgroup, represented by *P. maniculatus*, carries both active L1s and B1s, we know that B1 extinction happened after the split of the outgroup, yet the point at which B1 lost activity in the basal group is to be determined. Here we show that the peak of the most recent B1 family resides at ~11.3% in *O. palustris* and ~10.7% in *S. hispidus* (Table S2). These peaks reside in the same time window as L1 family OS2 (~11.1% in *O. palustris* and ~10.3% in *S. hispidus*, Table S1), suggesting that B1 family OS1 is coincident in time with L1 family OS2. Since L1 family OS2 is the youngest L1 family prior to the separation of the basal group and the L1-extinct clade, the last wave of B1 retrotransposition likely preceded the extinction of L1.

**Figure 5.**
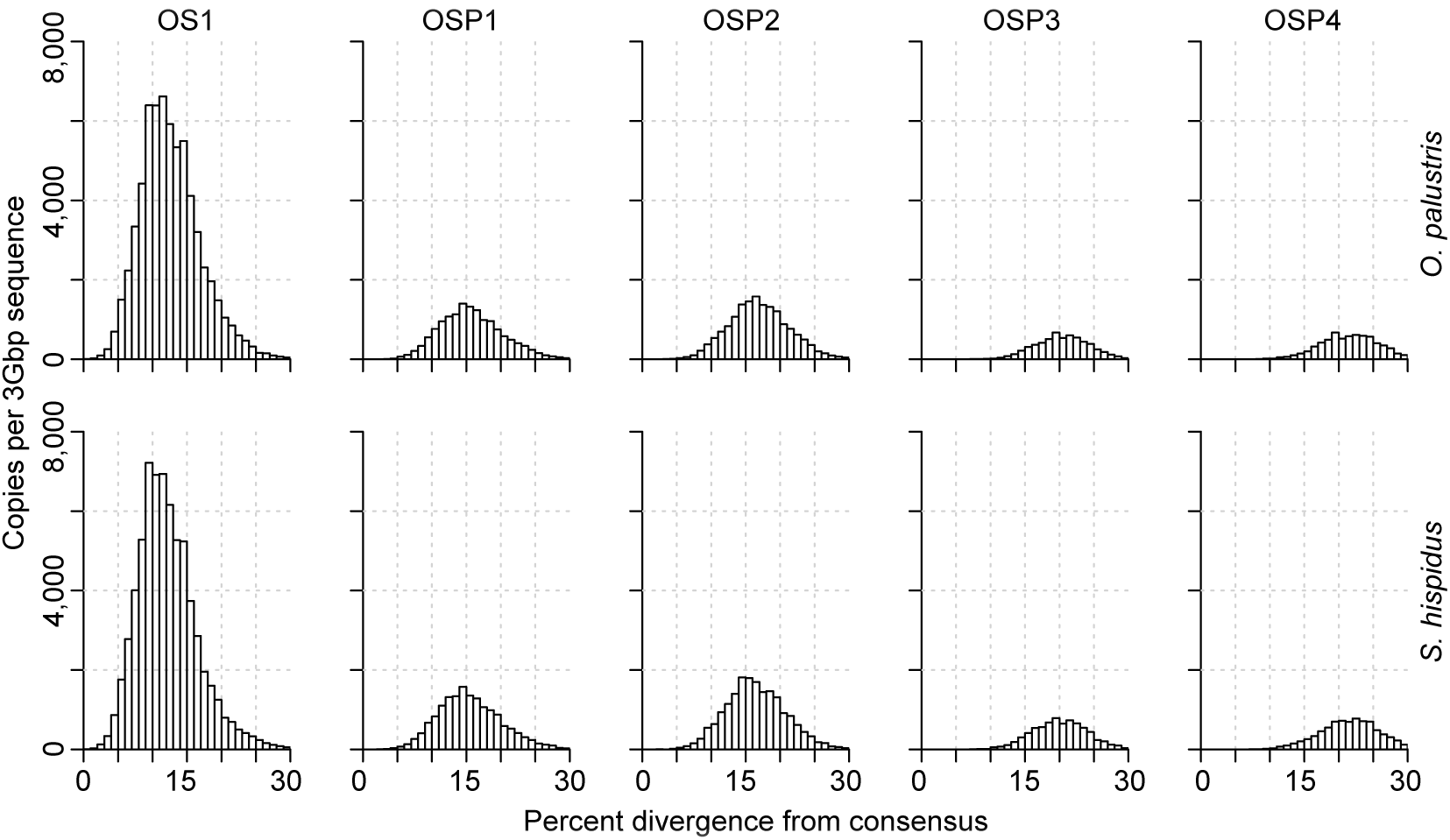
The age distribution of B1 families. B1 families in each row are arranged in chronological order with the youngest families on the left. The species analyzed in each row is indicated at the right. Names of families are noted on the top of each panel. B1 copy number is plotted by percent divergence from the corresponding subfamily consensus in 1% windows. The age of each family is approximated by the peak of the distribution. B1 copy numbers are normalized as copies per three Gbp of MiSeq sequence which approximates the copy number per haploid genome.

## Discussion

In this paper we explore the tempo of L1 and B1 activity surrounding the extinction of both elements that occurred in most species within the rodent subfamily Sigmodontinae. This work is made possible by sequencing methods that allow us to gather large amounts of sequence data and by the availability of a robust species phylogeny for the group (Figure 1). A recent phylogenetic analysis of muroid rodents [68] indicates that the tribe Sigmodontini is basal to the group and sister to the tribe Ichthyomyini. These two tribes are sister to a large, polytomic group (the Oryzomyalia) which includes the remaining five tribes; this group is the result of a rapid radiation of rodents into South America about 5 MYA [69]. Previous work indicated that L1s are extinct in the Oryzomyalia but active in the Sigmodontini, which includes one genus, *Sigmodon*, with 14 species. L1 extinction in the Oryzomyalia has been documented in 14 genera distributed across four tribes spanning this group (Figure 1) [56]. B1s are extinct in Oryzomyalia and Sigmodontini, but the status of both L1s and B1 in the intermediate tribe, Ichthyomyini, is unknown. Thus, L1 extinction from this single event likely affects between 345 and 362 species, or about 7% of all mammalian species.

We reconstructed the shared evolutionary history of L1s and B1s in Sigmodontinae in the period preceding and following extinction of these elements. Our results suggest that L1 master elements have been replaced steadily prior to the extinction of both L1 and B1. This is reflected by the consecutive series of L1 families shared by *O. palustris* and *S. hispidus* after their divergence from *Peromyscus*. B1 elements did not appear to take advantage of every wave of L1 activity, but a wave of L1 retrotransposition (family L1-OS2) corresponds to the B1 retrotransposition peak just prior to B1 extinction (B1-OS1).

There is reasonably strong evidence that L1 extinction occurred after the split between the L1-extinct clade and the basal group. A summary diagram showing the higher level of OS1 activity in *O. palustris* compared to *S. hispidus* (Figure 4A) suggests that the events leading to L1 extinction also happened after the split, rather than that a recovery occurred in *S. hispidus* as has been previously suggested [56]. The evolutionary history of B1 in *O. palustris* and *S. hispidus* is comparable. New B1 deposition into the genome was low except for the period directly preceding B1 extinction (Figures 4B and 5). Given the short length of B1s, it is more difficult to identify subfamily clusters, so our estimation of the timing of B1 extinction is weaker than for L1. However, two lines of evidence suggest that the last burst of B1 activity occurred prior to the split between the L1-extinct and basal groups. First, the peak activity of B1OSP1 corresponds most closely to the peak activity of L1OS2, which appears to precede the split of these two rodent clades. Secondly, there is no indication of large differences of activity for any of the B1 subfamilies, as was the case for L1. We suggest that finding the status of both L1s and B1s in the Ichthyomyini lineage might be critical to resolving the timing of B1 extinction.

The most challenging part of studying transposable element evolution history in rodents is the limitation of time windows reflected by detectable sequences. The sequences detectable by RepeatMasker decrease drastically beyond 40% divergence. Since the mutation rate in the rodent lineage is one of the highest in all mammals, 40% divergence in L1 and B1 traces back to the common ancestor of sigmodontine rodents and *P. maniculatus*, while similar studies on bats [54] and primates [70, 71] trace back to the common ancestor of mammals. Fortunately, *P. maniculatus* carries both active L1s and B1s and is close enough to serve as an outgroup in this study. We were able to identify an L1 family shared by *O. palustris*, *S. hispidus* and *P. maniculatus*, family OSP1.

However, there is an advantage of studying rodents in this type of evolutionary study. Since the mutation rate in the rodent lineage is higher than that of primates and bats due to shorter generation time, evolution in L1 and B1 families reflected by a given span of divergence covers a wider window of time compared to more slowly evolving species. This gives the age distributions of L1s and B1s higher resolution and allows us to discern subtle differences between subfamily ages.

This study is fully bioinformatics-based, but several points are important if one is to consider the underlying molecular events relevant to transpositional bursts and extinctions. L1 and B1 retrotransposition is regulated by a plethora of cellular factors [41-43, 52] and reliant on others [46, 47]. For evolutionary studies, especially the ones related to L1 and B1 extinction, the historical state of host cellular factors could dramatically change the retrotransposition landscape. Given that not all cellular factors that affect L1 and B1 retrotransposition are known and that coevolution between the elements and these cellular factors is expected, it is not currently possible to fully deduce the molecular events surrounding L1 extinction. However, from an evolutionary perspective, fixed retrotransposition events are recorded in the genome and evolve neutrally as pseudogenes unless excised or too old to be recognized. Therefore, the fossil record of L1s and B1s in the genome is a good temporal record of retrotransposition over time. However, one should keep in mind that estimation of retrotransposition rate based on historical L1 copy numbers could be affected by the excision rate of the host genome. It has been shown that the mammalian genomes have been constantly expelling sequences by various mechanisms and the excision rate varies in different clades of mammals [72]. As old insertions are not actively making new copies, they are exposed to the excision mechanisms for longer time, thus fewer copies of the older families are represented on the histogram. Old L1 and B1 copies also suffer from the recognition limitation of alignment algorithms. Detectable L1 and B1 copies are drastically reduced beyond 40% divergence.

## Methods

*O. palustris* and *S. hispidus* genomic DNA was sequenced in two separate batches using MiSeq (Illumina, Inc., San Diego, CA) at the IBEST Genomic Resources Core (University of Idaho, Moscow, ID). Paired-end libraries were generated with an insert size of 450-550 bp; ~13 and 14 million total reads were generated for *O. palustris* and *S. hispidus*, respectively. Sequences were processed with SeqyClean (https://bitbucket.org/izhbannikov/seqyclean) and the paired-ends were joined with FLASH [73]. Genome coverage was equivalent to approximately 1.5X; 5.47 Gbp of sequence were generated for *O. palustris* and 6.06 Gbp for *S. hispidus*, but we note that genome size within the sigmodontine rodents varies. Although the genome size of *O. palustris* is not documented to our knowledge, the genome size of sister species in *Oryzomys* suggests that *Sigmodon* genomes are 11-16% larger than those of *Oryzomys* [74].

L1 reconstruction for both species was generated based on partial genomic sequences generated by 454 Pyrosequencing (Roche Applied Science, Penzberg, Germany) at the IBEST Genomic Resources Core, 203 Mbp of sequence for *O. palustris* and 214 Mbp for *S. hispidus*. *P. maniculatus* genome trace files were obtained from NCBI. Reconstruction of the 3’ ends of *O. palustris* and *S. hispidus* L1s started with a 575 bp consensus seed in the 3’ half of L1 ORF2 generated following Cantrell *et al*. [75]. A bioinformatic pipeline for reconstructing a full length L1 is described by Yang *et al*. [54]. Briefly, sequences were acquired from the genome trace files based on percent identity. The overhangs of the found sequences allowed the creation of new seeds at both ends of the L1 fragment and were used to initiate another round of query. In this case, the reconstruction walk was repeated in the 3’ direction until the 3’ end of ORF2 was reached. Percent identity cutoff was set at 92% for *O. palustris* and higher percent identity (97 to 99%) was used for *S. hispidus* to assure a satisfactory consensus for each walk and the exclusion of older L1 elements. The 3’ 300 bp of the reconstructed L1s were then used as the reference sequences for COSEG analysis described below.

B1 sequences from Rinehart *et al*. [57] were used as starting seeds for B1 analysis. The PCR-amplified B1s from *O. palustris* and *S. hispidus* were aligned with Lasergene MegAlign (DNASTAR, Madison, WI) and the consensus sequence (146 bp) was used as the reference sequence for COSEG analysis.

L1 and B1 subfamilies in *O. palustris* and *S. hispidus* were identified and characterized in similar fashion as described below and are summarized in Table S1 and S2.

The reconstructed 300 bp sequences from the 3’ end of *O. palustris* and *S. hispidus* L1 ORF2 were each used as the initial L1 query sequences, and the full length B1 consensuses from each species, based on Rinehart *et al*. [57], were used as the initial B1 query sequences. *O. palustris* and *S. hispidus* MiSeq genomic DNA libraries were queried to identify homologous sequences using RepeatMasker [67] with default parameters. Hits from each search were filtered for >90% coverage of the query sequence and subsequently used for the first COSEG [67] (http://www.repeatmasker.org/COSEGDownload.html) run to identify subfamilies base on shared, co-segregating sequence variants. All COSEG runs were conducted under default parameter except as noted. Parameters were set such that at least 250 sequences were required to form an L1 subfamily and 1,000 were required to form a B1 subfamily. In order to identify older subfamilies, the consensus sequences of the subfamilies identified by the first COSEG run were used as queries to again search the *O. palustris* and *S. hispidus* MiSeq libraries using RepeatMasker. The identified sequences from the second RepeatMasker run were filtered for >90% coverage and extracted. *O. palustris* and *S. hispidus* sequences are combined and a second COSEG run was carried out on the combined sequences. To avoid the possible formation of random subfamilies due to the short length of B1 and the high copy number of the detected sequences, the sequences required to form a subfamily was increased from 1,000 (for the former separate run) to 2,000, whereas this number for L1 remained unchanged at 250. The consensus sequences of the resulting COSEG subfamilies were trimmed to exclude ends that were not common to all subfamilies and the CpG sites were removed and, thus, treated as gaps by RepeatMasker and not counted for the divergence calculation. These modified subfamily consensus sequences were used for a final query of the individual *O. palustris* and *S. hispidus* MiSeq libraries using RepeatMasker. Sequences from this third run were assigned to subfamilies based on percent divergence and this information was stored for further analysis.

*P. maniculatus* genome trace files were data-mined in a similar fashion through a single round of RepeatMasker and COSEG. The *O. palustris* L1 and B1 sequences described above were used as the initial query seeds for this run. Selected *P. maniculatus* subfamilies were used to demarcate the ages of the subfamilies identified in the *O. palustris* and *S. hispidus* genomes (Figure 3).

Subfamily consensus sequences generated by the second COSEG run of the *O. palustris* and *S. hispidus* libraries were combined and aligned with MegAlign using the Clustal W method for L1 or Clustal V method for B1 and a distance matrix was calculated based on the alignment. Based on the alignment, a maximum likelihood tree was constructed using PhyML [76] with the GTR+I+G model and 100 bootstrap replicates (Figure S1). L1 and B1 sequences were then assigned to families based on the topology of the tree and a no more than 3.5% within-family pairwise distance from their subfamily consensuses for L1 and 4.4% for B1. Given that the L1 and B1 masters are constantly being replaced during evolution, perfect designation of large families is not possible. The 3.5% threshold was chosen so as to cluster closely related subfamilies without inflating the number of families. Families are named according to their species-specificity and age: “S” indicates *Sigmodon*-specific families, “OS” for families shared by *Sigmodon* and *Oryzomys* and “OSP” for families shared by *Sigmodon*, *Oryzomys* and *Peromyscus*; numbers in family names indicates the age of a family within the family group with “1” being the youngest. Histograms of L1 and B1 age distributions were generated by R [77] histogram function using a window size of 1% (Figure 3). Percent divergence corresponding to retrotransposition peaks of individual families and subfamilies were determined by R using the kernel smoothing function with 0.4% bandwidth (Table S1 and S2).

## Availability of supporting data

All data generated or analyzed during this study are included in this published article and its supplementary information files.

## List of abbreviations

LINE: Long INterspersed Element
SINE: Short INterspersed Element
MYA: Million Years Ago
ORF: Open Reading Frame
*O. palustris*: *Oryzomys palustris*
*S. hispidus*: *Sigmodon hispidus*
*P. maniculatus*: *Peromyscus maniculatus*

## Author’s contributions

LY and HAW perceived and designed the experiment, analyzed the data and wrote the manuscript. LY prepared the DNA library for high-throughput sequencing and performed the bioinformatics analysis.

## Acknowledgements

We thank LuAnn Scott for helpful discussions, editing and proofreading of the manuscript. We thank Dr. Jerzy Jurka at the Genetic Information Research Institute for offering the bioinformatics training. We thank John Brunsfeld and Dr. Celeste Brown on helpful ideas of the L1 reconstruction pipeline design. We thank Drs. Wenfeng An, Celeste Brown and James Foster for helpful comments and discussions. We thank the IBEST Genomics Resources Core for helping us to generate the high-throughput sequencing data used and the IBEST Computer Resources Core for hosting the clusters used for the bioinformatics analysis. This work was funded by National Institute of Health R01-GM38737 to HW and National Science Foundation DDIG-1210694 to HW and LY; analytical resources were provided by National Institute of Health GM103324 and GM103408. The funder had no role in study design, data collection and analysis, decision to publish, or preparation of the manuscript.

## Supporting information

**Figure S1. The maximum likelihood phylogeny of detected L1 subfamilies.** Reconstructed *O. palustris* and *S. hispidus* L1s, labeled ‘seed’, and *P. maniculatus* subfamilies 5 and 6 are included as markers. The tree was reconstructed using PhyML [76] with the GTR+I+G model and 100 bootstrap replicates. Bootstrap values > 80% are shown.

**Figure S2. The age distribution of all detected L1 and B1 sequences.** Ages of sequences are approximated by their percent divergence from the corresponding subfamily consensus sequences and plotted in 1% windows. Species and retrotransposon names are indicated at the top of each panel.

**Table S1. The statistics and designation of L1 subfamilies and families.** “Ory” stands for *O. palustris* and “Sig” stands for *S. hispidus*. “Peak” indicates the peak of the L1 divergence distribution of the subfamily or family identified by kernel smoothing. Copy numbers are normalized as copies per three Gbp of MiSeq sequence used for the search, which approximates the copy number per haploid genome. Designation of families is only shown after the first subfamily that belongs to it; all subsequent subfamilies belong to this family until the demarcation of the next family. Characters in family names: “S” represents *S. hispidus*-specific, “OS” for shared by *O. palustris* and *S. hispidus* and “OSP” for shared by *O. palustris*, *S. hispidus* and *P. maniculatus*. Numbers in the family names reflect their ages among the family group with “1” being the youngest. Copy numbers of families are rounded sums of subfamily copy numbers per three Gbp of sequences and, thus, are occasionally off by one.

**Table S2. The statistics and designation of B1 subfamilies and families.** “Ory” stands for *O. palustris* and “Sig” stands for *S. hispidus*. “Peak” indicates the peak of the B1 divergence distribution of the subfamily or family identified by kernel smoothing. Copy numbers are normalized by per three Gbp of MiSeq sequence used for the search. Designation of families is only shown after the first subfamily that belongs to it; all subsequent subfamilies belong to this family until the demarcation of the next family. Characters in family names: “OS” represents families shared by *O. palustris* and *S. hispidus* and “OSP” for families shared by *O. palustris*, *S.hispidus* and *P. maniculatus*. Numbers in the family names reflect their ages within the family group with “1” being the youngest. Copy numbers of families are rounded sums of subfamily copy numbers per three Gbp of sequences.

